# The Computational Auditory Signal Processing and Perception Model (CASP): A Revised Version

**DOI:** 10.1101/2024.07.31.605576

**Authors:** Lily Cassandra Paulick, Helia Relaño-Iborra, Torsten Dau

## Abstract

This study introduces a revised version of the computational auditory signal processing and perception (CASP) model (Jepsen *et al*., 2008), which has undergone substantial updates and refinements aimed at enhancing its accuracy and usability across various applications. A primary motivation for the revision was the integration of a more realistic non-linear inner hair cell (IHC) model to address the limitations of its more simplistic predecessor. Demonstrating backwards compatibility, the revised model exhibited similar predictive power to previous CASP implementations across conditions of intensity discrimination, simultaneous and forward masking, and modulation detection, effectively accounting for data from normal-hearing listeners. Additionally, improved decision-making strategies and a streamlined model configuration further enhance the model’s usability and accessibility. Overall, the revised CASP offers a more accurate and intuitive framework for simulating auditory processing and perception across diverse conditions and tasks. The revised model may be a useful tool in studying the influence of the ear’s nonlinear response properties on internal representations, particularly concerning the effects of sensorineural hearing loss on auditory perception.

## I. INTRODUCTION

Computational auditory models offer mathematical frameworks for simulating how the auditory system processes sound, serving as powerful tools for exploring and comprehending its fundamental mechanisms (e.g.; Giguère *et al*., 1994; Heinz *et al*., 2001; Moore *et al*., 1988; Patterson and Moore, 1986; Zwicker and Scharf, 1965). Given the complexity and multi-faceted nature of auditory processing, diverse modeling approaches have emerged over the years, each tailored to specific applications and following different modeling philosophies. Some modeling approaches, commonly referred to as biophysical models (e.g.; Giguère *et al*., 1994; Lopez-Poveda and Eustaquio-Martín, 2006; Saremi and Stenfelt, 2013), aim to accurately replicate the underlying physiological mechanisms of the auditory system. These models are typically constructed based on detailed knowledge of the anatomy and physiology of the auditory system. In contrast, other approaches focus on predicting perceptual data. These perceptual models, also referred to as ‘functional’ or ‘phenomenological’ models, concentrate on reproducing specific psychoacoustic phenomena without explicitly modeling the precise biological mechanisms involved (e.g.; Dau *et al*., 1996a; Patterson *et al*., 1987). Instead, their aim is to replicate the observed ‘effective’ processing of a specific stage in the auditory pathway, such as modeling its input-output function. Ideally, they encompass a broad spectrum of phenomena within a simplified framework. While perceptual models excel at predicting outcomes of psychoacoustic experiments, their contribution to understanding underlying physiological processes is, by design, limited.

One example of such a perceptual model is the computational model of human auditory signal processing and perception (CASP; Jepsen and Dau, 2011; Jepsen *et al*., 2008). CASP serves as a functional model of the auditory periphery, addressing various aspects of simultaneous and non-simultaneous masking. Building on the success of the modulation filterbank model (Dau *et al*., 1997), Jepsen *et al*. (2008) enhanced the model to offer a more comprehensive description of peripheral processing. This was achieved by introducing outer and middle ear filters, along with significant modifications at the basilar membrane (BM) processing stage. Instead of utilizing a linear gammatone filterbank (Patterson *et al*., 1987) to simulate the frequency-selective processing of the BM, the CASP model adopted the dual-resonance non-linear filterbank (DRNL; Lopez-Poveda and Meddis, 2001). The DRNL filterbank not only replicated the BM’s frequency-selective behavior but also accounted for the level-dependent, non-linear cochlear filtering.

In addition to the outer and middle ear filters and the DRNL filterbank, CASP’s preprocessing stages include an inner hair cell (IHC) transduction stage, expansion, adaptation, and finally, a modulation filterbank. The inclusion of a non-linear BM processing stage expanded the model’s predictive capabilities to encompass tasks reflecting the BM’s level-dependent processing, such as spectral masking patterns obtained with different masker levels. The CASP model successfully predicted the behavior of listeners with normal hearing (referred to as ‘NH listeners’) across a wide range of psychoacoustic conditions, including intensity discrimination, spectral and temporal masking, and modulation detection (Jepsen *et al*., 2008). Furthermore, early versions of the framework have been utilized as a front-end for automatic speech recognition (Kleinschmidt *et al*., 2001; Tchorz and Kollmeier, 1999), speech quality assessment (Hansen and Kollmeier, 1999), as well as for the prediction of speech intelligibility (Relaño-Iborra *et al*., 2019).

In a subsequent work (Jepsen and Dau, 2011), the model was expanded to incorporate data from listeners with sensorineural hearing loss (HI listeners). This involved adjusting the nonlinear cochlear processing stage to account for individual estimates of the BM’s input-output function and introducing attenuation in the IHC model to simulate IHC loss. While successful in predicting average trends in listeners’ behavior in certain fundamental tasks (Jepsen and Dau, 2011; Jepsen *et al*., 2008), the framework notably underestimates the considerable variability observed in the data from HI listeners, especially in more complex speech-related tasks (Relaño-Iborra and Dau, 2022). These challenges may arise from several limitations, including simplifications in the signal processing stages represented in the model.

One potential factor limiting the predictive accuracy is the oversimplified IHC transduction stage of the CASP model. This stage converts the BM’s mechanical vibrations into receptor potentials within the IHCs through a process of half-wave rectification and low-pass filtering. Although this simplified approach captures the loss of phase-locking to the temporal fine structure of acoustic signals at higher frequencies, it fails to replicate the saturation of transduction at elevated sound pressure levels as documented in physiological studies (Dallos, 1985; Jia *et al*., 2007; Patuzzi and Sellick, 1983). This additional auditory compression stage may be crucial for encoding sounds of medium intensities, as indicated by recent studies (Carney, 2018; Carney *et al*., 2015). While the effect of IHC saturation on low-level signals is minimal, its asymmetrical non-linearity leads to notable differences in how signals at medium to high sound pressure levels are encoded. This saturation is especially pronounced around the characteristic frequency (CF), where it compresses the signal’s envelope tuned to the CF while processing frequencies distant from the CF in an (almost) linear fashion. Such variations across the auditory spectrum could be essential cues for describing a range of detection thresholds (Carney, 2018; Scheidiger *et al*., 2018).

The present study introduces and evaluates a refined CASP model, featuring an improved IHC transduction stage inspired by previous biophysical modeling efforts (Lopez-Poveda and Eustaquio-Martín, 2006; Shamma *et al*., 1986; Sumner *et al*., 2002; Verhulst *et al*., 2018). This advanced stage replaces the earlier simplified IHC transduction model, enabling a more thorough evaluation of the effects of the IHC saturation on internal stimulus representations and the anticipated perceptual effects.

Additionally, enhancements have been made to make the revised CASP model more intuitive and reduce its degrees of freedom. This addresses a limitation in the model’s backend, which uses an ideal observer concept based on matched-filtering (Dau *et al*., 1996a). In earlier CASP versions (Jepsen and Dau, 2011; Jepsen *et al*., 2008), experimenters had to manually define backend parameters, such as template level and the selection of relevant peripheral channels for specific detection tasks, based on their understanding of the task. This parametrization can significantly influence the model’s outcomes. In the revised model, an automated selection algorithm is introduced to eliminate the need for predefined input signal information for different psychoacoustic tasks. The algorithm automatically identifies the relevant region of the audio-frequency spectrum in response to the acoustic stimulus, setting up the model configuration without researcher intervention. The revised model also includes further enhancements in backend processing and parameter adjustments. The performance of the revised CASP is evaluated using data from NH listeners in various psychoacoustic conditions.

## II. THE REVISED CASP MODEL

The CASP model comprises two primary components as outlined below: the preprocessing stage, which involves the processing steps that create the internal representations of the input signal along the modeled auditory pathway, and the backend processing stage, which examines the internal representation(s) to assess the detectability of a particular signal. The structure of the revised CASP model is illustrated in Figure 1. Panel A provides a description of the preprocessing stages, while Panel B illustrates the model’s decision backend. The input to the model is an acoustic signal (measured in Pascals), which undergoes processing through filters that simulate the outer and middle ear functions (Lopez-Poveda and Meddis, 2001), followed by a DRNL filterbank (Lopez-Poveda and Meddis, 2001) that captures non-linear, level-dependent frequency selectivity of the peripheral auditory system. The resulting frequency-decomposed signal is then passed through the updated IHC transduction stage. This revised version employs a biophysically informed non-linear IHC model, replacing the previously simplified version. Adaptation effects are modeled with a series of feedback loops (Püschel, 1988), and the signal is further processed through a modulation filterbank to mimic frequency selectivity in the modulation envelope domain. The output is a three-dimensional internal representation that includes time, audio frequency, and modulation frequency dimensions. This internal representation can be visualized as a series of ‘modulation spectrograms’ (Relaño-Iborra *et al*., 2019) as illustrated in Figure 1. These modulation spectrograms are then analyzed in the model’s backend (Figure 1, Panel B), utilizing the ideal observer method (Dau *et al*., 1996a) to evaluate the detectability of stimuli, particularly suited for alternative-forced-choice (AFC) tasks. The model’s resolution is constrained by constant-variance internal noise, a feature consistent with previous versions of the model (Dau *et al*., 1996a; Jepsen *et al*., 2008), where this constant noise variance is set during a calibration condition. Additional details about both the preprocessing and backend stages, including the updates made in the revised CASP model, are elaborated below. A parameter set is given in Appendix A.

**FIG. 1.**
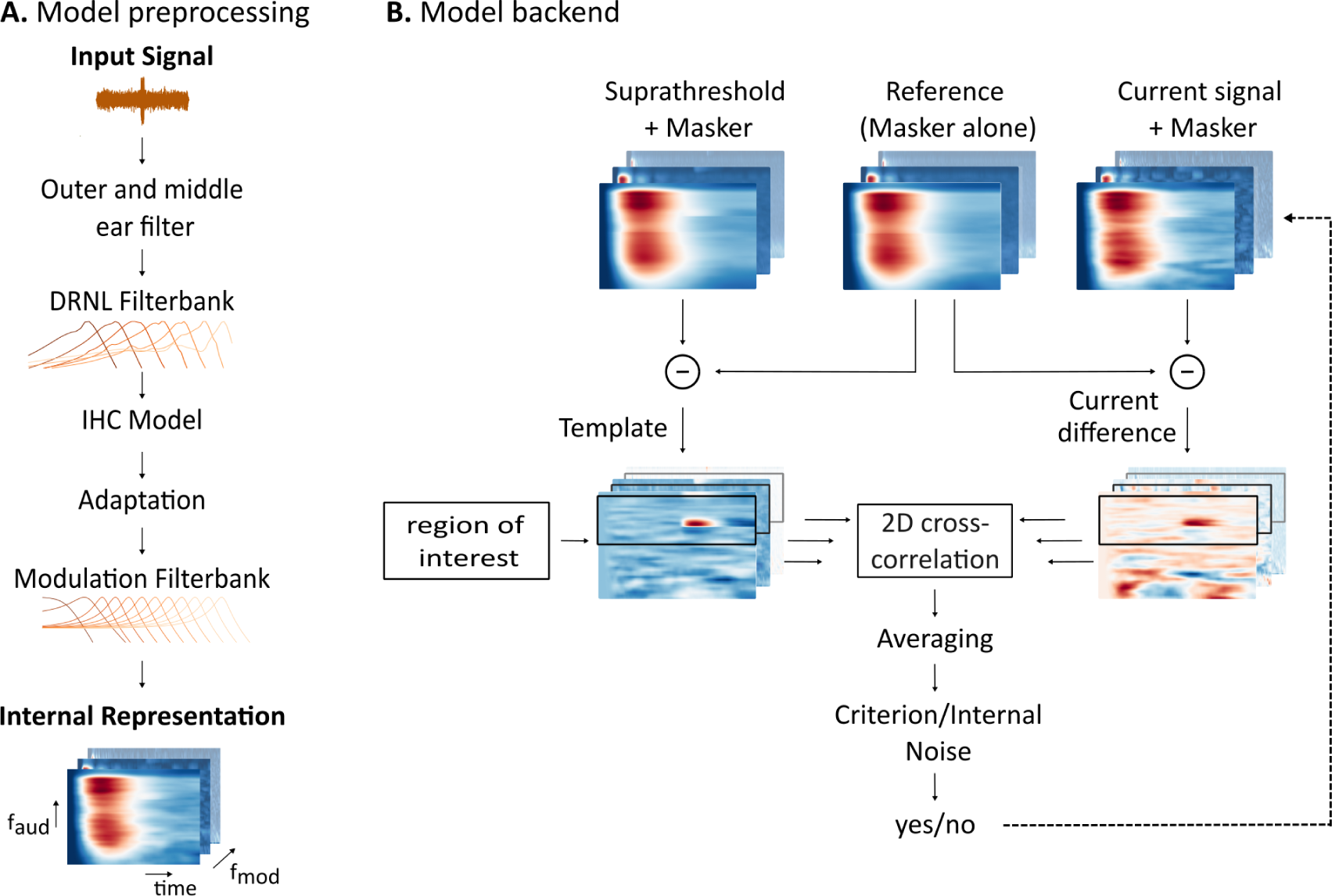
The CASP model comprises two distinct components: (A.) preprocessing stages responsible for constructing the internal representation from an acoustic input signal, and (B) the backend which utilizes an optimal detector to generate a decision metric using the internal representations of a template, reference, and target signal.

### A. Outer- and middle-ear filter and DRNL filterbank

In the initial stage of the model, the acoustic signal (in Pascals) is processed through an outer- and middle ear filter, implemented as a pair of linear-phase finite impulse response filters (Lopez-Poveda and Meddis, 2001). The output from this stage approximates the peak velocity of the stapes in the middle ear, which then feeds into the DRNL filterbank. The DRNL filterbank consists of a set of filters that process the signal along two parallel paths: one linear and one compressive non-linear path (Meddis *et al*., 2001). The linear path includes a linear amplification step, followed by a series of gammatone filters and then a series of low-pass filters. The non-linear path processes the signal through gammatone filters, applies a broken-stick non-linear function, and then passes the signal through additional gammatone and low-pass filters. The outputs from both paths are combined, with the non-linear path being more influential at low to medium sound pressure levels (SPLs), and the linear path dominating at higher SPLs, typically above ∼ 70 dB SPL. This setup effectively simulates the level-dependent compression and auditory filter tuning characteristics of the BM. The filters within this model are configured to cover 50 frequencies, equally spaced on an equivalent rectangular bandwidth scale (Glasberg and Moore, 1990), ranging from 250 to 8000 Hz, although the actual range of auditory frequencies is restricted in the backend of the model as described further in section II F. The DRNL’s parameterization from the original model has been largely maintained, with minor adjustments detailed in the appendix. More comprehensive information on these stages is provided in Lopez-Poveda and Meddis (2001) and Jepsen *et al*. (2008).

### B. IHC model

The revised IHC model closely aligns with previous biophysical modeling studies (Altoè *et al*., 2017; Lopez-Poveda and Eustaquio-Martín, 2006; Shamma *et al*., 1986; Sumner *et al*., 2002) that based their characterizations of IHC transduction on animal data (Dallos, 1985, 1986; Jia *et al*., 2007; Johnson *et al*., 2011; Russell *et al*., 1986). The implementation included in the revised CASP takes the BM motion from the DRNL filterbank stage as its input. The BM motion is then transformed into stereocilia displacement *u*(*t*) using a scaling constant similar to the approach used by Verhulst *et al*. (2018). The cilia displacement modulates the apical conductance *G*_MET_(*u*), which is modeled by a second-order Boltzmann function (Altoè *et al*., 2017; Kros *et al*., 1992; Sumner *et al*., 2002):

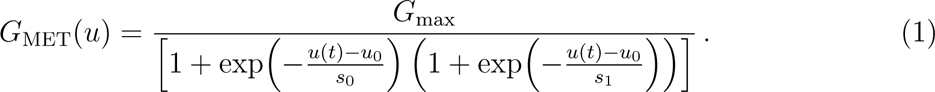

Here, *G*_max_ represents the maximum mechanical conductance, and *u*_0_*, s*_0_ and *s*_1_ are parameters that define the shape of the compressive curve. These parameters were selected in line with the IHC model used by Verhulst *et al*. (2018), informed by recent in vivo studies by Jia *et al*. (2007) and Johnson *et al*. (2011). The IHC receptor potential is then modeled using a passive electrical circuit analogy (Altoè *et al*., 2017; Sumner *et al*., 2002), represented by the differential equation:

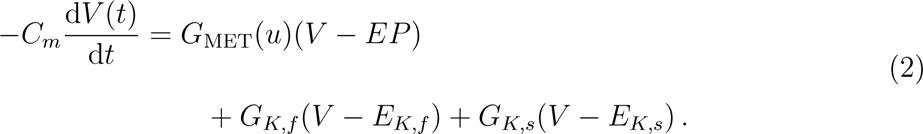

Here, *C_m_* is the cell’s capacitance, *EP* represents the endocochlear potential, *G_k,f/s_* are the fast and slow potassium conductances, and *E_K,f/s_* the corresponding reversal potentials. For simplicity and to reduce the number of parameters, the model assumes that the potassium (*K*^+^) currents are voltage-independent (Shamma *et al*., 1986; Sumner *et al*., 2002), diverging from models that assume voltage dependency (Lopez-Poveda and Eustaquio-Martín, 2006; Verhulst *et al*., 2018). Specific values for all relevant parameters are provided in Appendix A. This approach successfully captures key properties of the IHC transduction process: the loss of phase-locking to the temporal fine structure at higher frequencies (Palmer and Russell, 1986; Russell and Sellick, 1983) and the saturation effect in IHC transduction (Dallos, 1986; Kros *et al*., 1990). The new model introduces a more nuanced representation of IHC transduction, substituting the previous implementation (Dau *et al*., 1996a; Jepsen *et al*., 2008) that applied a half-wave rectification and low-pass filtering with a cut-off frequency of *f*_cut-off_ = 1 kHz to the input signal. Following this, the previous model required a squaring expansion stage that converted the signal into an intensity-like representation (Jepsen *et al*., 2008). While the low-pass filter accounted for the observed loss of phase-locking to the temporal fine structure at higher frequencies (Palmer and Russell, 1986; Russell and Sellick, 1983), the half-wave rectification process was inadequate in representing non-linear, saturating characteristics of the IHC’s input-output function.

Figure 2 illustrates the internal representations generated by both the original CASP model with the linear IHC transduction stage (CASP) and the revised CASP model that includes a non-linear IHC stage (CASP_rev_), in response to two different stimulus types. To demonstrate the loss of phase-locking at higher frequencies, Panel A shows the response at the output of the IHC model within the on-frequency channel to pure tones with frequencies ranging from 300 Hz to 5 kHz. These tones, 200 ms in duration with 20 ms hannning windows applied, were presented at 70 dB SPL. For both versions of the model, there is a noticeable reduction in the alternating current (AC) component of the signals as the frequency increases, indicating a loss of temporal fine structure. However, the degree of remaining fine structure at higher frequencies differs between the models, with the linear IHC model exhibiting a more substantial reduction.

**FIG. 2.**
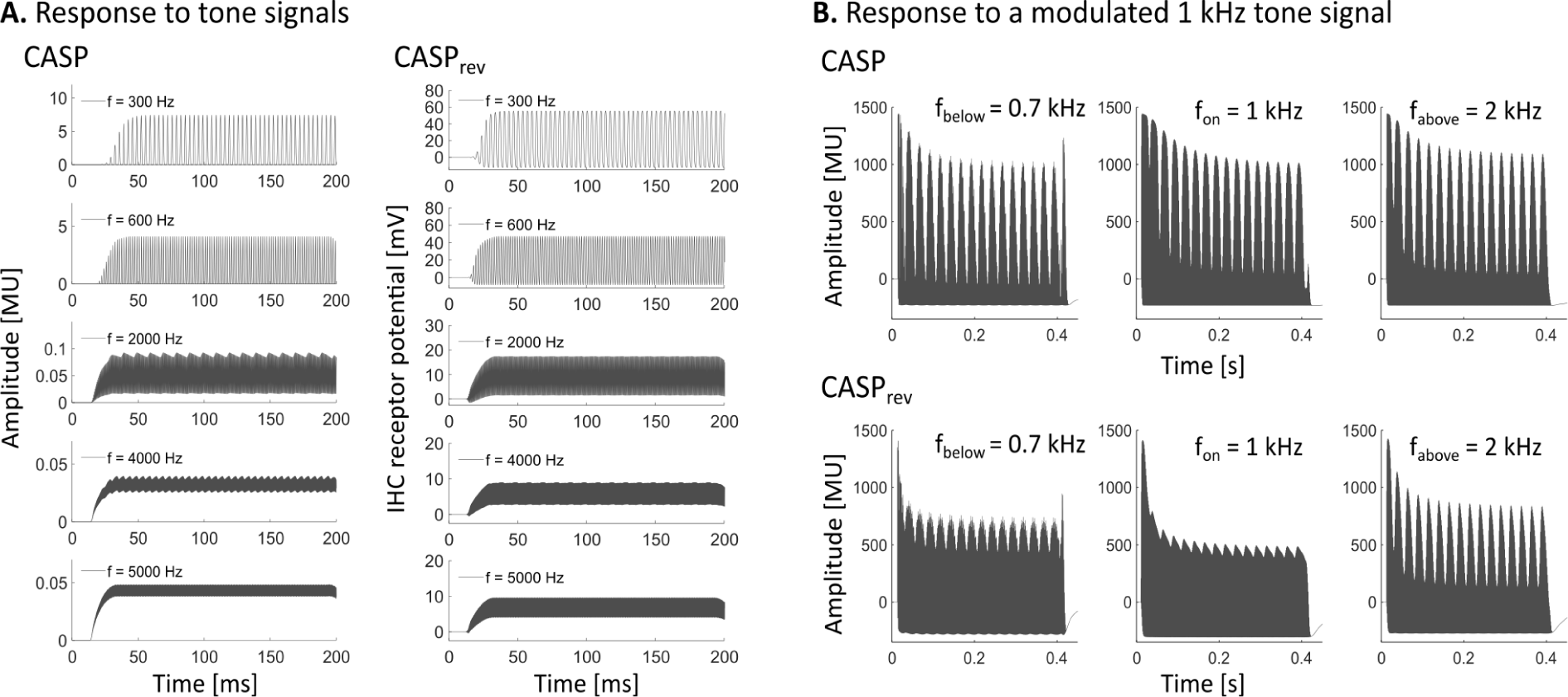
Panel A shows internal representations at the output of the IHC stage in response to tones at various frequencies at the the on-frequency channel. The left column shows simulations from the original CASP with the linear IHC model (CASP) while the right column presents outcomes from the revised CASP that includes the non-linear IHC model (CASP_rev_). Panel B shows internal representations at the output of the adaptation stage, generated in response to a 40 Hz-modulated 2 kHz tone, for both the CASP model with the linear IHC stage (top) and the non-linear IHC stage (bottom). The internal representations are depicted at peripheral channels below CF (0.7 kHz, left), at CF (1 kHz, middle) and above CF (2 kHz, right).

To illustrate the effects of the IHC saturation, Panel B in Figure 2 shows the model responses to a 1 kHz tone at 85 dB SPL, modulated with a 40 Hz rate and modulation depth of −5 dB. The internal representations are shown at the output of the adaptation stage (described in section II C) for the on-frequency channel (1 kHz), as well as for channels below and above the on-frequency channel (tuned to 0.7 kHz and 2 kHz, respectively). While both models include compression effects at the BM level, the non-linear IHC model introduces additional compression in the on-frequency responses (middle panels). This IHC saturation in the non-linear IHC model leads to a ‘flattened’ response in the on-frequency channel, differing from the internal representation produced by the linear model. Furthermore, CASP_rev_ shows a more pronounced difference across frequency channels in the responses, as signals are processed linearly away from the CF, resulting in distinct processing for on- and off-frequency channels. This new behavior is consistent with physiological data that showed IHC responses to several tone frequencies (Dallos, 1985; Russell and Sellick, 1983).

### C. Adaptation

The adaptation stage aims to capture adaptive properties of the auditory periphery (Smith, 1977; Westerman and Smith, 1984) by employing a series of feedback loops (Püschel, 1988), each with a distinct time constant. Together, these loops apply an approximately logarithmic compression to stationary input signals while preserving a more linear transformation for inputs with rapid changes. In earlier versions of the model (Dau *et al*., 1996a), originally designed for linearly processed inputs, the time constants of the feedback loops had been adjusted to fit results from NH listeners in a forward masking task. In the present study, forward masking simulations demonstrated the need to adjust these parameters to accomodate the non-linearly processed inputs from the revised IHC stage. The time constants were modified to exponentially distributed values from 0.007 to 0.5 s, specifically *τ* = [0.007, 0.032, 0.088, 0.214, 0.5] s to fit forward masked thresholds. This modification moderately diminishes the sharp onset effects associated with the shorter time constants. Furthermore, the minimum input level for the adaptation stage was tailored to each channel, based on the International Organization for Standardization (2019) threshold levels for the characteristic frequency at the stage’s input. This allows the model to more accurately predict hearing thresholds in quiet. Lastly, the onset response was maximally limited to ten times that of the steady state response. Further details on the adaptation stage can be found in Dau *et al*. (1996a).

### D. Modulation processing

The outputs from the adaptation stage are further decomposed into modulation subbands using a first-order low-pass filter (*f*_cut-off_ = 150 Hz) followed by a modulation filterbank (Dau *et al*., 1997). This modulation filterbank comprises a sequence of frequency-shifted resonance filters, functioning as band-pass filters, specifically tuned to modulation frequencies spanning from 2.5 to 1000 Hz. Filters tuned to modulation frequencies below 10 Hz are assigned a constant bandwidth of 5 Hz while those above 10 Hz adopt a logarithmic scaling with a constant *Q* factor of 2. Furthermore, to simulate the loss of envelope phase sensitivity at higher modulation frequencies (Dau *et al*., 1997, 1996b; Langner and Schreiner, 1988), for filters centered above or at 10 Hz the absolute values of the modulation filter outputs are taken, attenuated by a factor of ✓2, whereas for filters centered below 10 Hz the real part is used. For each peripheral channel, the modulation sub-bands are confined to those with a center frequency below one-quarter of the center frequency of the respective peripheral channel (Langner and Schreiner, 1988; Verhey *et al*., 1999).

### E. Backend processing

The preprocessing stage of the model generates a three-dimensional internal representation of the input signal, encompassing time, audio frequency and modulation frequency dimensions. These internal representations are subsequently analyzed in the backend processing, which closely follows the ideal observer rationale of the original model (Dau *et al*., 1996a). The decision process is designed to simulate human behavior in an adaptive alternative-forced-choice (AFC) procedure. At the beginning of each experimental run, the model establishes a stored reference signal derived from the average of several internal representations of the masker-alone signal. Additionally, it forms a stored template from the normalized difference between the average of several internal representations of the supra-threshold-plus-masker signal and the reference (see Figure 1, B). In each trial, the current interval signals have the reference subtracted and are subsequently compared to the template. This comparison employs a non-normalized 2D-correlation across the time and audio frequency dimensions, represented by indices *t* and *n*, respectively, as follows:

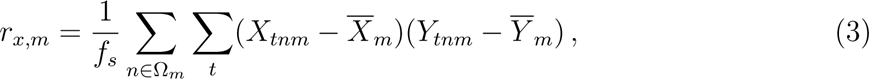

where *f_s_* refers to the sampling frequency, *Y* = [*T* × *N* × *M*] denotes the three-dimensional template, with *T* referring to the length of the signal, *N* the number of auditory channels and *M* the number of modulation channels. 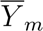 represents the mean across both the time and frequency dimensions, and *X* = [*T* ×*N* ×*M*] the current difference signal and its corresponding mean 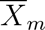. Here, the range of audio frequency channels considered (the subset Ω*_m_*) is restricted to those selected by the channel selection algorithm described below (section II F) and satisfying the condition of being above four times the given modulation frequency *f_m_*. For each interval *X*, the outcome is a correlation value *r_m_* for each modulation frequency, which are subsequently added to produce a single correlation value *r_x_*. The model classifies a signal as detectable if the difference in correlation between the template and the target interval and that of the template and noise intervals is higher than the resolution limit, defined by a constant internal noise variance under a fitting condition (Dau *et al*., 1996a). Similar to previous model implementations, the internal noise variance is calibrated once to satisfy Weber’s law in an intensity discrimination task.

### F. Channel selection

In the revised model, an automated algorithm is introduced to identify a region of interest, which includes relevant peripheral channels to be incorporated into the decision metric, thereby determining the precise model configuration. The algorithm analyses the template (section II E) at the level of the adaptation stage in order to make a selection in the auditory channel domain. The frequency channels of this internal representation are processed through a moving average filter to smooth out noisy variations or fluctuations and reveal underlying patterns. The window size for the moving mean analysis is adapted to each center frequency, corresponding to *τ* = 25*/f_c_*, ensuring that the smoothing is appropriately matched to the frequency. After smoothing, the root-mean-square (rms) is calculated for each frequency channel, and the maximum is found. Subsequently, frequency channels with an rms above one half of the maximum are identified, and the region of interest is defined from the lowest to the highest frequency channel that satisfies this condition, including all the channels in between. Channels outside this range are discarded and thus do not contribute further to the model’s decision metric. Additionally, a constraint is applied to ensure that the region of interest spans at least half one octave below to one octave above the maximum. The channel selection is performed once per experimental run, implying that the region of interest is kept constant across the adaptive AFC procedure for each experimental parameter. This algorithm is introduced to overcome a limitation of previous CASP implementations, where the simulation operator was required to preselect the relevant peripheral channels (i.e., model configuration) before each experimental run. This selection depended on the experimenter’s prior understanding of the specific psychoacoustic task and their expertise. In contrast, this algorithm reduces reliance on preexisting information and decreases uncertainty levels.

## III. METHODS

The revised CASP was tested under three experimental paradigms reflecting intensity coding, temporal acuity, and modulation processing. Human data of NH listeners on intensity discrimination (Miller, 1947; Viemeister and Bacon, 1988), forward masking (Jepsen *et al*., 2008), and modulation detection Dau *et al*. (1997) from previous studies were simulated. For each psychoacoustic task, the procedure and stimulus details are provided below. In addition, the supra-threshold level for the template generation is specified, along with the relevant frequency range used for simulations with the original CASP configuration, i.e., without the automatic selection algorithm. In all cases, an adaptive AFC procedure was employed using the AFC-Toolbox 1.40 for MATLAB (Ewert, 2013), utilizing an n-AFC paradigm with an adaptive tracking rule mirroring the procedures conducted with human listeners. For each experiment, the results were averaged over five runs of the adaptive procedure.

### A. Notes on the original CASP implementation

The performance of the revised model is compared to both the human data and predictions from the original model. Notably, the original model predictions do not directly correspond to the simulations presented in the original publication by Jepsen *et al*. (2008). Despite considerable effort, reproducing these simulations proved challenging due to the limited description of the parameter set and implementation details provided in the paper. Therefore, this paper implements the model as accurately as possible using the parameters provided in the original publication. Simulations using the original model structure (i.e. without IHC saturation) could potentially be improved by adjusting the parameter set. However, we chose to use the parameters reported in the original paper to maintain consistency. The primary focus of the present paper is to validate the revised implementation, rather than to optimise the original model’s performance.

### B. Calibration

To determine the constant variance of the internal noise in the optimal detector, a fitting condition is initially carried out. This noise variance represents the resolution limitation of the model (see, section II E). An intensity discrimination task at 60 dB is selected for this purpose. The model is calibrated to satisfy the observed just-noticeable difference (JND) of approximately 0.47 dB at a 60 dB presentation level for a 1 kHz tone (Viemeister and Bacon, 1988) and a JND of 0.44 dB for a broadband noise signal from 150 Hz to 7 kHz (Miller, 1947). In this setting, the tone had a duration of 200 ms with raised cosine ramps of 10 ms applied at both the onset and offset, while the noise lasted for 1.5 s with 50 ms raised-cosine ramps applied. The determined variance remained unchanged for other measurement points and across all subsequent psychoacoustic tasks.

### C. Intensity discrimination

An intensity discrimination task was simulated for a 1 kHz tone and broadband noise spanning from 150 Hz to 7 kHz. To compare to data from Viemeister and Bacon (1988) for the tone and Miller (1947) for the noise, the just noticeable difference (JND) was measured at 20, 30, 40, 50, 60, and 70 dB SPL and 20, 25, 35, 45, 55, and 70 dB SPL, respectively. Note that the 60 dB point constituted the fitting condition, and thus - unlike the other points - does not represent a true model prediction. The tone had a duration of 200 ms with raised cosine ramps of 10 ms applied at the onset and offset. As in the reference condition, a 2-AFC with 1-up-2-down tracking rule that tracks the 70.7% point on the psychometric function was used. The noise lasted for 1.5 s with 50 ms raised-cosine. Here, a 3-AFC with 1-up-1-down rule is used to find the JNDs of the model. The supra-threshold template level was set to 5 dB above the test signal level and averaged across 15 representations. For the original CASP model, which requires manual selection of the region of interest, the relevant channels were confined to +*/*− an octave below and above the target frequency in the case of the tone. In the case of the broadband noise, frequency channels ranging from 100 Hz to 8 kHz were included, as reported in Jepsen *et al*. (2008).

### D. Forward masking

To evaluate aspects of temporal resolution in the model, a forward masking task with noise maskers was simulated, following the procedure in Jepsen *et al*. (2008). Here, a 3-AFC with 1-up-2-down tracking rule that tracks the 70.7% point on the psychometric function was used. Threshold functions were determined for the detection of a 4 kHz tone played after broadband noise at three different levels: 40, 60 and 80 dB SPL. The noise masker was band-limited Gaussian noise, spanning frequencies from 20 to 8000 Hz and with a duration of 200 ms, along with 2 ms raised-cosine ramps applied. The tone signal had a duration of 10 ms with a Hanning window applied over the entire signal duration (Jepsen *et al*., 2008).

Thresholds were obtained for temporal separations between the masker offset and signal onset of −20, −10, −5, 0, 5, 10, 20, 40, 80 and 150 ms. Human data from Jepsen *et al*. (2008) were used for comparison. The supra-threshold level for the template was set to 10 dB above the masker level and averaged over 15 representations. For the original model configuration without channel selection, the predetermined frequency range was limited to 3.6 to 5 kHz (Jepsen *et al*., 2008).

### E. Modulation detection

In the modulation detection task, narrowband Gaussian noise centered at 5 kHz was utilized as a carrier. The carrier bandwidths were set at 3, 31 and 314 Hz. The carriers were presented at 65 dB SPL with a duration of 1 s, incorporating 200 ms raised-cosine ramps. Sinusoidal amplitude modulation with modulation frequencies of 3, 5, 7, 10, 20, 30, 50 and 100 Hz was applied to modulated the carriers. The tracking variable was the sinusoidal modulation depth expressed on a dB scale as *M* = 20 ∗ log(*m*), where *m* represents the modulation depth ranging from 0 to 1. A 3-AFC with 1-up-2-down tracking rule that tracks the 70.7% point on the psychometric function was used. This procedure mirrors that of Dau *et al*. (1997), whose human data were used for comparison. The supra-threshold level for the template was set to a modulation depth of −6 dB and averaged across 15 representations. In previous simulations using the original model configuration only the on-frequency channel tuned to 5 kHz was considered (Jepsen *et al*., 2008). However, our simulations with the original model implementation revealed that using only the on-frequency channel was insufficient to obtain thresholds in the given task. Therefore, the range was expanded to include channels from 4.5 to 8 kHz.

## IV. RESULTS

Model predictions were compared to the corresponding human data (black) taken from the literature. For each task, two sets of model predictions are represented: one from the original CASP model using the linear IHC model (CASP, red) and the other from the revised CASP model using the non-linear IHC model (CASP_rev_, blue), with shaded areas indicating standard deviations. As described in section III, the frequency range for the original model is predetermined as in Jepsen *et al*. (2008), whereas the revised model employs the proposed automatic channel selection algorithm, as described in section II F. The accuracy of the models is evaluated in terms of their Pearson’s correlations (r) with the data, as well as the mean average errors (MAE) between the predictions and the measurements.

### A. Intensity discrimination

Intensity discrimination tasks asses a listener’s ability to discern changes in stimulus intensities. It is often quantified in terms of the just noticeable difference (JND) in stimulus intensity, a measure commonly governed by Weber’s law which suggests an approximately constant JND across different stimulus levels (Miller, 1947). As in the original model, the constant variance internal noise combined with the approximately logarithmic compression in the adaptation loops yields a roughly constant JND across all stimulus levels.

Figure 3 shows the model predictions alongside the reference data (Miller, 1947; Viemeister and Bacon, 1988) for the 1 kHz tone (Panel A) and the broadband noise (Panel B). In the tone intensity discrimination tasks, a slight decrease in the JND can be observed towards higher signal levels, commonly referred to as the near miss to Weber’s law. This decrease is commonly attributed to the spread of excitation to channels tuned above the tone frequency at elevated sound pressure levels. The revised CASP model shows a slight increase in JND towards lower sound pressure levels, exhibiting a mean absolute error (MAE) of 0.27 dB. Conversely, the original model shows a constant JND for levels from 20 to 60 dB, and a slight decrease in JND at 70 dB. The overall MAE is 0.23 dB. Notably, the model estimations rely both on the DRNL filterbank to simulate the spread of excitation with increasing level and the adaptation loops that apply an approximately logarithmic compression.

**FIG. 3.**
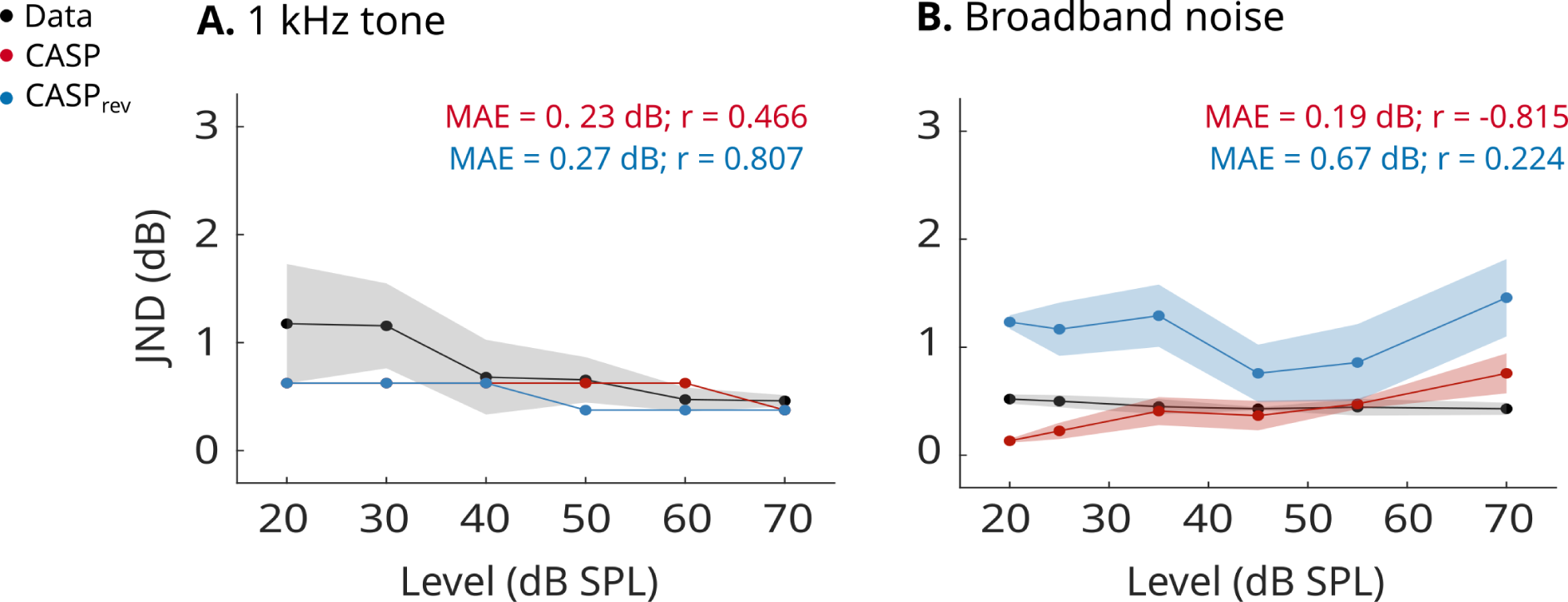
The just noticeable difference (JND) in stimulus intensity is shown as a function of level for a 1 kHz tone (Panel A) and broadband noise (Panel B). Human data (Miller, 1947; Viemeister and Bacon, 1988) is shown in black alongside model predictions with the original CASP model (red) and the revised model (blue). For each model prediction, the mean absolute error (MAE) and Pearson’s correlation value (r) are reported.

For the broadband noise, the revised model predicts a on average 0.67 dB larger JND compared to the data. On the other hand, the original model displays a slight, steady increase in JND with rising levels. Here, the model slightly underestimates the JND by approximately 0.39 dB at the lowest level and overestimates it by approximately 0.33 dB at the highest sound pressure level.

### B. Forward masking

Figure 4 shows the masking thresholds of the 4 kHz tone for the three masker levels (Panel A. 40 dB, Panel B. 60 dB, Panel C. 80 dB). Human data are represented in black, while model predictions are depicted in red and blue for the original and revised model, respectively, with shaded areas indicating standard deviations. The data indicate a faster decrease in threshold with smaller onset-offset intervals, gradually slowing as delays increase. Across all masker levels, the thresholds converge to a level of approximately 12 dB at a delay of 150 ms, mirroring the threshold of the 10 ms long tone. Negative onset-offset intervals represent simultaneous masking conditions, where the tone is embedded within the noise.

**FIG. 4.**
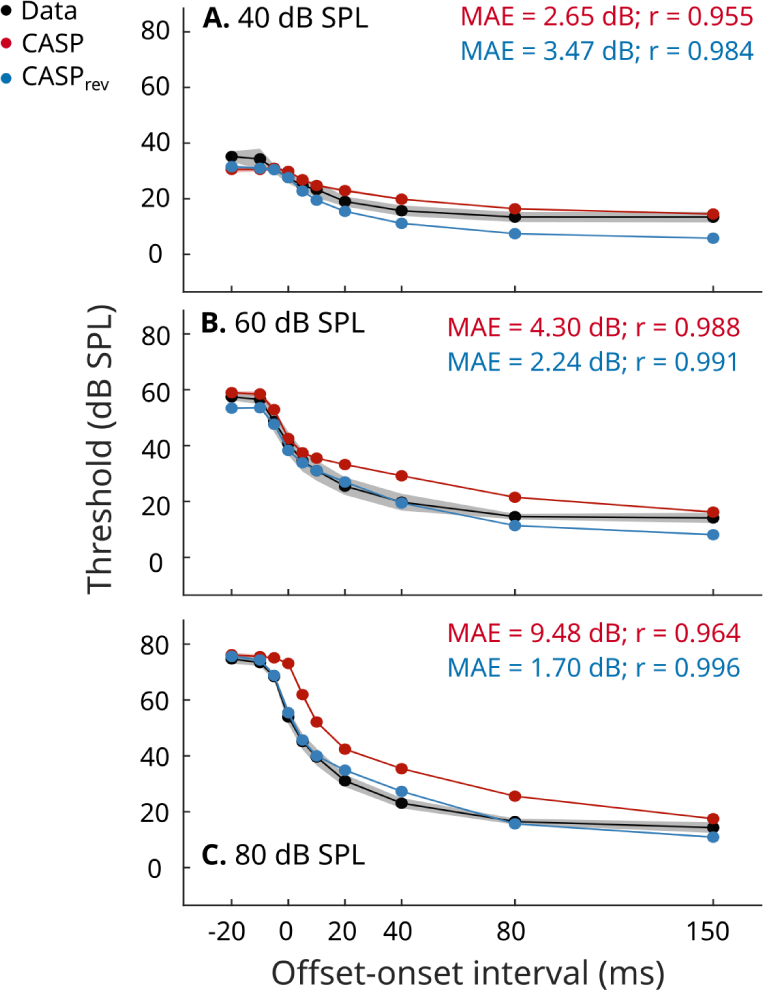
Masked threshold curves as a function of the onset-offset interval between masker and signal are shown for a forward masking paradigm with three different masker levels, 40 dB (Panel A), 60 dB (Panel B), 80 dB (Panel C). The target signal was a 10 ms long 4 kHz tone. Measured data from Jepsen *et al*. (2008) is indicated in black and model predictions for the original and revised CASP in red and blue, respectively. For each model prediction, the mean absolute error (MAE) and Pearson’s correlation value (r) are reported.

Both models are able to capture these trends, with the revised model generally exhibiting more accurate performance in the given metrics. Particularly, the revised model closely aligns with the data for all three masker levels, albeit underestimating the thresholds slightly in the 40 dB masker level case towards larger delays. The original model displays a poorer performance, and generally overestimates threshold levels in all three cases, especially towards delays above 20 ms.

### C. Modulation detection

In the modulation detection task, temporal modulation transfer functions (TMTFs) for narrow-band noise carriers were simulated and compared to data from Dau *et al*. (1997). The noise carriers, centered at 5 kHz, were generated with three different bandwidths (3, 31 and 314 Hz), each resulting in distinct TMTF patterns due to the inherent statistical fluctuations of the narrow-band carriers. Figure 5 illustrates the modulation depth at threshold, in dB, for the reference data (black) and two model configurations (CASP, in red; CASP_rev_, in blue). Overall, both models exhibit similar performance and effectively capture the distinct patters across the different bandwidths. For the narrowest carrier (3 Hz wide), both models considerably underestimate thresholds at higher modulation frequencies by around 7 dB. For the 31-Hz carrier the high-pass characteristic is observable in the model predictions, but the thresholds are overestimated slightly by both model configurations. In the 314 Hz wide carrier case, both models display a good match with the human data above 10 Hz, but show an increase in threshold towards the small modulation frequencies that is not reflected in the data. This is more pronounced in the predictions from the revised model.

**FIG. 5.**
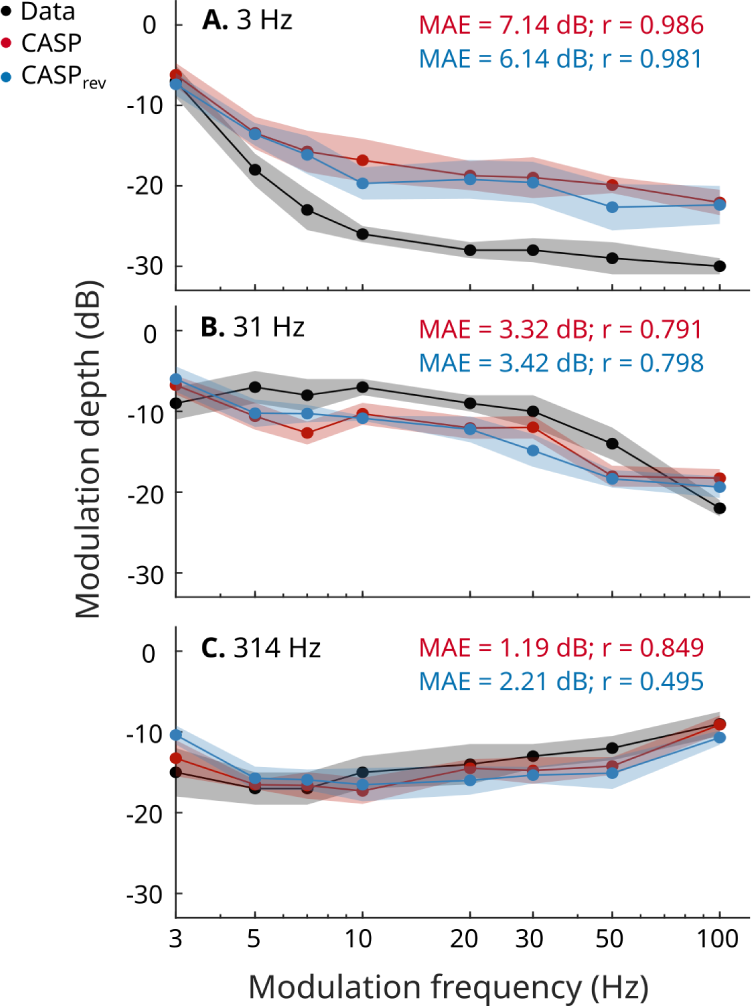
Temporal modulation transfer functions (TMTFs) are shown for amplitude modulated noise carriers, centred at 5 kHz, with three different bandwidths: 3 Hz (Panel A.), 31 Hz (Panel B.) and 314 Hz (Panel C.). Measured data (Dau *et al*., 1997) (black) is shown alongside simulated TMTFs of the original CASP model (red) and the revised CASP model (blue). For each model prediction, the mean absolute error (MAE) and Pearson’s correlation value (r) are reported.

## V. DISCUSSION

This study presented the revised CASP, which underwent multiple updates and refinements across several stages, aiming to enhance its accuracy and usability for various applications. Across the simulated conditions presented here, the revised model demonstrated backwards compatibility, exhibiting similar predictive power to previous CASP implementations.

A key motivation for the revised model was the integration of a more realistic non-linear IHC model to address the inherent limitations of the simplistic IHC representation in its predecessor. Incorporating an additional non-linear model stage introduces challenges, as non-linear stages can interact in complex ways. This requires a detailed understanding of the associated transformations. While the different model stages are computationally independent, each processing stage influences the operating point and dynamic range of the successive stages, necessitating ongoing adjustments along the signal path. Moreover, non-linear models often require the addition of new modeling parameters, demanding careful tuning constrained by the availability of valuable experimental data. This challenge was already exemplified with the transition from a linear gammatone filterbank to the more sophisticated DRNL filterbank (Jepsen *et al*., 2008). This transition required both the addition of outer- and middle-ear filters to produce realistic inputs to the BM model and an additional squaring device before the adaptation stage to account for forward masking. In the revised model, the more realistic IHC model obviates the need for this expansion stage. However, it required an adjustment of the time constants and minimum input levels to the adaptation stage, which were originally designed for linearly processed inputs.

Despite these complexities, the present study demonstrated that including IHC non-linearities in the overall model is feasible without compromising its established predictive power across various conditions of simultaneous and non-simultaneous masking. In tasks such as intensity discrimination, forward masking, and modulation detection, the revised CASP model performed comparably to the original CASP model, effectively accounting for data from NH listeners. Moreover, conditions such as spectral masking patterns, as presented in Jepsen *et al*. (2008), are anticipated to remain largely unaffected by the IHC non-linearity, given that the detection cues here are principally driven by the DRNL stage.

The goal of the revised CASP extended beyond merely enhancing predictive accuracy; it aimed to enhance realism in the processing pathway and reduce reliance on external fitting. The comparable performance of the two models prompts reflection on the necessity of incorporating IHC saturation for the chosen tasks and highlights the trade-off between model complexity, realism, and predictive accuracy. While realism is a crucial goal, it does not always result in significant improvements in predictive power, as demonstrated by the findings presented here. Hence, it is essential to consider the increase in computational complexity and its predictive benefits within the broader context of the model’s overarching goal.

Moreover, incorporating realistic processing stages enhances the model’s potential for generalization across a broader range of tasks and conditions. The discernible impact on model performance may have been mitigated by the complex interplay of multiple model components and the already largely accurate predictions in the presented conditions. For tasks heavily reliant on specific processing stages, such as the effects of the adaptation stage on forward masking, differences across other stages in the model configurations may be less pronounced. However, greater disparities are anticipated in more complex auditory tasks or studies involving listeners with hearing impairment, which have revealed limitations and challenges in individualizing the CASP model (Relaño-Iborra and Dau, 2022).

As illustrated in Figure 2, the non-linear IHC model yields a more distinct contrast and realistic signal representation across frequency channels. This outcome arises from the disparate processing pathways at the CF and away from CF. Consequently, the revised model’s performance is greatly influenced by the selection of peripheral channels for the decision metric, underscoring the significance of automatic selection over manual decision-making. Specifically, in the modulation detection task, both compression from the BM and the IHC stage contribute to reduced sensitivity to amplitude modulations in the on-frequency channel, where the response is effectively ‘flattened’ at the given level. Therefore, the revised CASP model may inaccurately predict TMTFs when relying solely on the on-frequency channel (centred at 5 kHz), as done in previous simulation setups of the model (Dau *et al*., 1997; Jepsen *et al*., 2008). Unlike these previous versions, which achieved accurate predictions with a single-channel approach due to less realistic processing, the revised model suggests that relevant cues for modulation detection exist away from the on-frequency channel, where modulations are more strongly encoded due to the more linear processing pathways. The discrepancy between simulations reported in Jepsen *et al*. (2008) using only the on-frequency channel and our simulations with the original model configuration, which failed to obtain thresholds with only the on-frequency channels, may indicate that the region of compression in the DRNL filterbank is shifted. This shift would suggest that the signal was processed more linearly in the former case, effectively increasing the model’s sensitivity to modulation. Although the two presented models here yield comparable predictions, they may employ different decision-making strategies, with distinct frequency channels dominating the correlation metric. Further investigations into the model’s performance in modulation detection tasks at various presentation levels (representing regions of linear and non-linear processing) and its impairment, leading to an effective compression loss, could provide insights into the role of IHC saturation in modulation detection tasks.

A crucial advancement in the revised CASP is the shift from manually-determining backend parameters to implementing an automated selection algorithm. This transition streamlines the model’s configuration, making it more universally applicable across various psychoacoustic tasks by reducing the need for task-specific parameter adjustments. This improvement facilitates result replication, enhancing user-friendliness and accessibility for interested researchers.

## VI. CONCLUSIONS

This study introduced a revised computational model of auditory processing and perception, aiming to overcome limitations of its predecessor by incorporating more realistic preprocessing stages and enhancing backend processing. Through validation across a range of psychoacoustic conditions for NH listeners, the findings demonstrated that integrating an additional non-linear processing stage at the level of the IHC does not compromise the model’s predictive power.

Further investigations could explore the applicability and adaptability of the revised CASP model to individuals with sensorineural hearing loss. Previous versions of the model adjusted the DRNL filterbank to accommodate individual basilar-membrane input-output functions, and introduced an attenuation after the IHC stage to simulate IHC loss. The inclusion of more realistic processing stages and added non-linearity has the potential to improve model predictions for this population, addressing some of the variability in the observed data that the original model struggled to capture (Jepsen and Dau, 2011). Additionally, evaluating the revised processing stages integrated within the speech-based sCASP model could provide valuable insights into how both NH and HI listeners process more complex signals.

## ACKNOWLEDGMENTS

This work was carried out in connection to the Center for Applied Hearing Research (CAHR) supported by Widex, Oticon, GN ReSound, and the Technical University of Denmark.

## APPENDIX A: MODEL PARAMETERS

In the following table, model parameters for the different stages of the revised model are presented. A full model implementation can be found at mu https://gitlab.com/lpau/casp_forafc.git.

**TABLE I.** Model Parameters.

|  |  |  |  |
| --- | --- | --- | --- |
| **DRNL filterbank** |  |  |  |
| *Linear pathway:* |  |  |  |
| $N_{GT,lin}$ | Number of cascaded GT filters | 2 | |
| $N_{LP,lin}$ | Number of cascaded LP filters | 4 | |
| | $\log_{10}(parameter) = p_0 + m \log_{10}(CF)$ | $p_0$ | $m$ |
| $g$ | Linear gain | 4.20405 | -0.47909 |
| $BW_{lin}$ | Bandwidth of GT filters | 0.03728 | 0.75 |
| $CF_{lin}$ | Characteristic frequency of GT filters | -0.06762 | 1.01 |
| $LP_{lin,cutoff}$ | Cutoff frequency of LP filters | -0.06762 | 1.017 |
| *Non-linear pathway:* |  |  |  |
| $N_{GT,nlin}$ | Number of cascaded GT filters | 2 | |
| $N_{LP,nlin}$ | Number of cascaded LP filters | 1 | |
| | | $p_0$ | $m$ |
| $a$ | Broken stick non-linearity parameter | 1.40298 | 0.81916 |
| $b$ | Broken stick non-linearity parameter | 1.61912 | -0.81867 |
| $c$ | Broken stick non-linearity parameter | -0.60206 | 0 |
| $BW_{nlin}$ | Bandwidth of GT filters | 0.03193 | 0.77 |
| $CF_{nlin}$ | Characteristic frequency of GT filters | -0.05252 | 1.0165 |
| $LP_{nlin,cutoff}$ | Cutoff frequency of LPfilters | -0.05252 | 1.0165 |
| **IHC model** |  |  |  |
| $c_{BM,cilia}$ | cilia/BM coupling gain | $5.62 * 10^{-6}$ | |
| $G_{max}$ | maximum mechanical conductance | 30 nS | |
| $u_0, s_0, s_1$ | Boltzmann function parameters | 20, 16, 35 nm | |
| $C_m$ | cell capacitance | 12.5 pF | |
| $EP$ | endocochlear potential | 90 mV | |
| $G_{K,f/s}$ | potassium conductance (fast/slow) | 19.8 nS | |
| $E_{K,f}$ | potassium reversal potential, fast | 71 mV | |
| $E_{K,s}$ | potassium reversal potential, slow | 78 mV | |
| **Adaptation loops** |  |  |  |
| $\tau_n$ | time constants | [0.007, 0.032, 0.088, 0.214, 0.5] s | |
| limit | overshoot limitation | 10 |  |
| $lvl_{min}$ | minimum input level | CF-specific vector | |
| **Modulation filterbank** |  |  |  |
| $LP_{cutoff}$ | 1st order lowpass filter cut-off frequency | 150 Hz | |
| BW | Bandwidth for $f_m = 5$ and $f_m = 10$ Hz | 5 Hz | |
| Q | Q-factor for $f_m > 10$ Hz | 2 | |
| **Optimal Detector** |  |  |  |
| $\sigma$ | internal noise variance | 40 | |

